# A functional role for the epigenetic regulator ING1 in activity-induced gene expression in primary cortical neurons

**DOI:** 10.1101/149450

**Authors:** Laura J. Leighton, Qiongyi Zhao, Xiang Li, Chuanyang Dai, Paul R. Marshall, Sha Liu, Yi Wang, Esmi L. Zajaczkowski, Nitin Khandelwal, Arvind Kumar, Timothy W. Bredy, Wei Wei

## Abstract

Epigenetic regulation of activity-induced gene expression involves multiple levels of molecular interaction, including histone and DNA modifications, as well as mechanisms of DNA repair. Here we demonstrate that the genome-wide deposition of Inhibitor of growth family member 1 (ING1), which is a central epigenetic regulatory protein, is dynamically regulated in response to activity in primary cortical neurons. ING1 knockdown leads to decreased expression of genes related to synaptic plasticity, including the regulatory subunit of calcineurin, *Ppp3r1*. In addition, ING1 binding at a site upstream of the transcription start site (TSS) of *Ppp3r1* depends on yet another group of neuroepigenetic regulatory proteins, the Piwi-like family, which are also involved in DNA repair. These findings provide new insight into a novel mode of activity-induced gene expression, which involves the interaction between different epigenetic regulatory mechanisms traditionally associated with gene repression and DNA repair.

**Author contributions:** L.J.L., Q.Z., T.W.B and W.W. designed the experiments. N.K., A.K., X.L., C.D., S.L. and W.W. designed and assembled shRNA constructs. L.J.L., W.W., X.L., C.D., P.R.M., E.Z., and S.L. conducted experiments. Q.Z. and Y.W. analysed ChIP-seq data. L.J.L., Q.Z., and W.W. wrote the paper. All authors reviewed and edited the manuscript.

**Conflicts of interest:** None.

## Introduction

Epigenetic mechanisms, including DNA methylation, ATP-dependent chromatin remodelling factors, post-transcriptional modification of histone proteins, and various non-coding RNA pathways, are now recognised as fundamental for the regulation of activity-induced neuronal gene expression. While some of these epigenetic pathways are relatively well known, others remain poorly characterised, and ongoing research continues to reveal novel epigenetic regulators in post-mitotic neurons that were previously believed to respond only to stress, DNA damage, or replication error. One such mechanism is the recent discovery that DNA double-strand breaks are induced in an experience-dependent manner, and associated with transcriptional activity (Suberbielle *et al.*, 2013; Bunch *et al.*, 2015; Madabhushi *et al.*, 2015); however, the precise mechanism by which this process interacts with canonical epigenetic mechanisms remains to be determined.

Inhibitor of growth family member 1 (ING1), initially characterised as a tumour suppressor gene involved in cell cycle arrest and apoptosis and associated with several forms of cancer (Nouman *et al.*, 2003; Ythier *et al.*, 2008), has recently been identified as a central epigenetic regulator. ING1 acts as both a transcriptional activator and repressor through interactions with DNA and a multitude of epigenetic regulatory proteins, including histone modifications, by modulating the activity of the mSIN3A/HDAC1 transcriptional repression complex (Skowyra *et al.*, 2001; Binda *et al.*, 2008). It is also a reader for the histone mark H3K4me3, and drives active DNA demethylation at H3K4me3 sites by recruiting the DNA damage response factor GADD45*α* (Schäfer *et al.*, 2013). In addition, ING1 contributes to transcriptional regulation of the miRNA-processing gene *Dgcr8*, and loss of ING1 has been shown to lead to the dysregulation of miRNA expression in mouse fibroblasts (Gómez-Cabello *et al.*, 2010). Finally, there is substantial evidence for a fundamental role for ING1 in DNA repair. ING1 is upregulated in response to DNA damage (Cheung Jr *et al.*, 2000; Cheung *et al.*, 2001; Ceruti *et al.*, 2013) and mutant forms of ING1 derived from tumours are unable to promote DNA repair (Ceruti *et al.*, 2013). Moreover, overexpression of ING1 enhances the repair of UV-damaged DNA in a p53-dependent manner (Cheung *et al.*, 2001). Whether ING1 is also involved in DNA repair in differentiated neurons, and represents an underlying mechanism of activity-induced gene expression, is currently unknown.

Another epigenetic pathway that is also involved in DNA repair and transcriptional regulation is the Piwi pathway. Piwi-like proteins are an evolutionarily conserved subset of the Argonaute family with similarity to *Drosophila* PIWI (Juliano *et al.*, 2011; Iwasaki *et al.*, 2015), and are required for the biogenesis and activity of Piwi-interacting RNAs (piRNAs). piRNAs comprise a distinct class of small non-coding RNAs which have several characteristic features, including occurring in genomic clusters, a tendency to have uracil as the first base, a length of 26-31nt, and 2’-O-methylation on the terminal nucleotide (Iwasaki *et al.*, 2015). To date, it is thought that the dominant function of piRNAs is to protect the genome by controlling the activity of transposons (Juliano *et al.*, 2011; Iwasaki *et al.*, 2015), and although they have a broad role in stem cell maintenance and reproduction in many species, their activity in mammals occurs predominantly within the male germline (Iwasaki *et al.*, 2015). As indicated, Piwi proteins have been linked to DNA repair; Yin *et al.* have reported that *Piwil2* lies upstream of the DNA damage response pathway, and is also activated by DNA damage in fibroblasts. Preventing *Piwil2* activation blocks histone H3 acetylation and subsequent chromatin relaxation that is necessary for DNA repair, resulting in a DNA repair defect (Yin *et al.*, 2011). Furthermore, the Piwi pathway has recently been implicated in neuronal function and brain organisation (Rajasethupathy *et al.*, 2012; Zhao *et al.*, 2015; Nandi *et al.*, 2016; Viljetic *et al.*, 2017) with numerous reports of piRNA expression in the mammalian brain (Lee *et al.*, 2011; Saxena *et al.*, 2012; Ghosheh *et al.*, 2016; Nandi *et al.*, 2016).

Given the potential overlap between ING1 and the Piwi pathway as epigenetic regulators, we investigated whether ING1 is involved in activity-induced gene expression and whether Piwi plays a role in this process. To model neuronal activation, we cultured mouse primary cortical neurons and used potassium chloride to induce depolarisation. This well-established tissue culture model recapitulates many of the cellular and transcriptional programs that occur in neurons in response to physiological stimulation occurring *in vivo* (Flavell *et al.;* Greer & Greenberg, 2008; Kim *et al.*, 2010; Ataman *et al.*, 2016).

Our findings indicate that ING1 is required for activity-dependent regulation of several genes relevant to neuronal plasticity and function, and that the function of ING1 in regulating a specific target gene, *Ppp3r1*, can be modulated by the Piwi pathway. This is the first evidence of an interaction between these two conserved epigenetic pathways, revealing a novel mode of activity-induced epigenetic regulation of gene expression in primary cortical neurons.

## Experimental Procedures

### Adeno-associated virus construction and packaging

For knockdown of *Piwil1* and *Piwil2*, pAAV2-mu6-shRNAs were constructed as described previously (Hommel *et al.*, 2003), with slight modifications. In brief, siRNAs were designed using *siDirect* version 2.0 (Naito *et al.*, 2004), which were synthesised along with their antisense strand separated by a loop sequence to form a hairpin. Forward and reverse strands of shRNAs were annealed and cloned into the pAAV-MCS vector, containing the mU6 promoter and eGFP, in between Xbal and Sapl restriction sites.

HEK293T cells were grown to 80% confluence in 150mm dishes. Lipofectamine was used to transfect cells with the plasmids AAV1, AAV2, pFdelta6, and pAAV-MCS. Transfected cells were cultured for a further 4 days, then collected and lysed by 3 freeze-thaw cycles. The cell lysate was filtered, treated with Benzonase and AAV particles were concentrated by ultracentrifugation. Concentrated viral pellets were resuspended in PBS and snap-frozen. Three AAVs were produced, one carrying a scrambled control shRNA with no specificity to any known mouse transcript and one targeting each of *Piwil1* and *Piwil2.* The combination of packaging plasmids AAV1 and AAV2 in a 1:1 ratio produced approximately a 1:1 mixture of AAV serotypes 1 and 2, which have a complementary transduction pattern in neurons.

### Lentivirus construction and packaging

Lentivirus was packaged according to our previously published protocol (Lin *et al.*, 2011). Briefly, HEK293T cells were grown to 70% confluence in triple-layer flasks. Lipofectamine was used to transfect cells with the plasmids pMDG, pRSV-rev and pMDLg/pRRE and the transfer vector (gene-specific shRNA cloned into FG12). Transfected cells were cultured for 48 hours, after which the culture medium was collected, clarified, filtered, and lentivirus particles concentrated by ultracentrifugation. Concentrated viral pellets were resuspended in PBS and snap-frozen. Four lentivirus were produced: one carrying a control shRNA with no specificity to any known mouse transcript, and the other three targeting ING1 (ING1 shRNA 1, 2, 3).

### Culture of primary mouse cortical neurons

Experiments involving animals were approved by the Animal Ethics Committee of the University of Queensland. Pregnant female C57BL/6 mice were euthanized at E16 and embryos immediately collected into ice-cold PBS. Embryonic cortices were dissected and enzymatically dissociated with 5U/mL papain at 37°C for 20 minutes. Papain was washed out with neuronal culture medium (Neurobasal medium supplemented with 1% GlutaMAX and 2% B-27). The cells were then mechanically dissociated, counted, and plated at a density of 750-1000 cells/mm^2^ in culture dishes coated with poly-D-ornithine. Neuronal cultures were maintained in an incubator at 37°C with 5% CO2. The culture medium was replaced 24-48 hours after preparation of the cultures, and thereafter cultures were fed every third day by replacement of 50% of the culture medium. To knock down expression of genes of interest, cultured neurons at 2-3 days in vitro were exposed to lentivirus or AAV for 12 hours, followed by replacement of the culture medium. To investigate activity-dependent gene regulation, cultured neurons at 7-10 days in vitro were depolarised by the addition of 20mM KCl to the culture medium for three hours. The cells were collected immediately for molecular analysis following KCl exposure.

### Culture of primary mouse embryonic fibroblasts

Pregnant female C57BL/6 mice were euthanised at E16 and embryos immediately collected into ice-cold PBS. Embryos were decapitated and the thoracic and abdominal organs discarded; remaining tissue was minced and stored at 4°C overnight in 3 volumes of 0.05% trypsin. Following incubation at 37°C for 45 minutes, tissue was washed and cells were mechanically dissociated, counted, and plated into uncoated T75 flasks at 25% density in Dulbecco’s modified Eagle medium (DMEM) supplemented with 10% fetal bovine serum. Cells were maintained in an incubator at 37°C with 5% CO2; cultures were passaged 3 times before use, at which time cultures were free from cells of non-fibroblast morphology.

To induce ING1 activity, MEFs at passage 3 were treated with ultraviolet light to damage genomic DNA. Growth medium was removed and cells washed once with Hank’s buffered salt solution, then placed without liquid into a Stratalinker (Stratagene) and irradiated with 40 J/m^2^ (0.87 J/m^2^/s at 254nm). Following irradiation, growth medium was replaced and cells were returned to the incubator for approximately 1-3 hours before collection for molecular analysis.

### Chromatin immunoprecipitation

Chromatin immunoprecipitation (ChIP) was performed according to our previously published protocol (Wei *et al.*, 2012). Briefly, neurons were fixed in 1% formaldehyde to establish protein-DNA crosslinks. Crosslinked neuronal lysates were sheared by sonication in ChIP lysis buffer to generate chromatin fragments with an average length of 300-500 bp. Immunoprecipitation was carried out using an antibody specific to ING1 (Cell Signaling Technologies #14625) overnight at 4°C; antibody-protein-DNA complexes were isolated by incubation with Protein G Dynabeads (Thermo) for 1h at 4°C, followed by three washes in low-salt buffer, and three washes in high-salt buffer. The precipitated protein-DNA complexes were resuspended in ChIP dilution buffer, then incubated for 4 hours at 65°C to reverse formaldehyde cross-linking. Protein was removed by Proteinase K digestion and the ING1-associated DNA fragments were recovered by phenol-chloroform extraction and ethanol precipitation.

### ChIP-Seq and sequencing data analysis

ING1-associated DNA isolated by ChIP was used to generate paired-end (PE) sequencing libraries following the protocol of the Illumina DNA kit. Libraries were sequenced using the Illumina HiSeq 2500 sequencing platform, with the read length of 126bp*2. Image processing and sequence data extraction were performed using the standard Illumina Genome Analyzer software and CASAVA (v1.8.2) software. Cutadapt (v1.8.1) was used to cut the adaptor sequences as well as low quality nucleotides at both ends (with the option of “-q 20,20”). Reads were aligned to the mouse reference genome (mm9) using BWA (v0.6.2) (Li & Durbin, 2009). Samtools (v0.1.17) (Li *et al.*, 2009) was then used to convert “SAM” files to “BAM” files, sort and index the “BAM” files, and remove duplicate reads. Reads with low mapping quality (<20) or which were not properly paired-end aligned to the reference genome were excluded from the downstream peak calling analysis. MACS (v1.4.2) (Zhang *et al.*, 2008) was used to call peaks for each sample with the parameter setting “-f BAM –keep-dup=all –nomodel –shiftsize 100 –g mm –p 1e-5 –bdg”. Peak summits identified by MACS from all samples were collected to generate a list of potential binding sites. Peak summits located within 600bp of each other were grouped together using a custom PERL script as these nearby peaks may represent the same binding site. Peaks that were detected by MACS in at least 2 biological replicates of either KCl+ or KCl− samples were defined as potential ING1 binding sites and only these sites were used for downstream analysis in this study. We then categorised all potential binding sites into six groups, including promoter (defined as 2kb upstream of transcription start sites in known genes), 5’ UTR, CDS, intron, 3’ UTR, and intergenic. To determine the ING1 occupancy between KCl+ and KCl− samples, a custom PERL script was applied to count the number of fragments (hereafter referred to as ‘counts’) that cover the peak summit in each sample. Each pair of properly aligned PE reads covering the peak summit represents one count. The total counts in each sample were normalised to 10 million for the normalised coverage plot between conditions.

### RNA isolation and reverse transcription

To extract RNA, cells were lysed with Nucleozol (Macherey-Nagel) directly in the culture dish and RNA was isolated following the manufacturer’s instructions. Reverse transcription was carried out using the QuantiTect kit (QIAGEN) using the provided RT Primer Mix and following the manufacturer’s instructions.

### Quantitative PCR for ChIP samples

Quantitative PCR reactions were prepared in duplicate, using 2x SYBR master mix (Qiagen), 500μM of each primer, and 1μL of template DNA purified as described previously after chromatin immunoprecipitation. Reactions were run on the Rotor-Gene Q platform and results were analysed using the delta-delta-CT method normalised to the input sample. Data was analysed using Student’s t-test or one-way ANOVA, as appropriate.

### Quantitative PCR for gene expression

Quantitative PCR reactions were prepared in duplicate, in a 10μL reaction volume, using 2X SYBR master mix (Qiagen), 500μM of each primer, and 1μL per reaction of a cDNA sample (the cDNA dilution factor varied according to target abundance). Reactions were run on the Rotor-Gene Q platform and results were analysed using the delta-delta-CT method, normalised to the reference gene *Pgk* (phosphoglycerate kinase). Data was analysed using Student’s t-test or one-way ANOVA as appropriate.

## Results

### Neural activity leads to a pronounced change in the genome-wide deposition of ING1

To determine whether ING1 functions as an activity-dependent epigenetic regulator in neurons, we first validated an antibody (Cell Signaling Technologies #14625) for chromatin immunoprecipitation (ChIP). ING1 ChIP was performed on mouse embryonic fibroblasts with and without ING1 induction via UV irradiation, and qPCR detected successful enrichment of the known ING1 target *Mageb2* (Schäfer *et al.*, 2013) from the UV-treated MEFs using this antibody (Figure 1a).

We next performed ChIP-seq, using the same antibody, to examine the genome-wide distribution of ING1 in primary cortical neurons in response to KCl-induced depolarisation (n=3 per group). A total of 275 binding sites for ING1 were identified and found to be predominantly localised to intergenic regions (69.09% KCl+/−) (Figure 1b). Of the 275 ING1 binding sites, 116 sites (42.18%) showed specific KCl-induced binding (Figure 1c). Therefore, although the global distribution of ING1 binding did not change in terms of broad genomic regions where ING1 is deposited, the qualitative interaction with the genome was altered, with significantly more and different genomic loci targeted following KCl-induced depolarisation (Figure 1b and 1c; Appendix A). In response to neuronal stimulation, ING1 binding was predominantly located (~53%) within 10kb upstream of the TSS of the nearest gene, representing a regulatory region including the promoter and potentially a cis-acting enhancer. Another 39% of ING1 binding sites overlapped with coding genes (including binding sites in the exon, intron, 5’ or 3’ UTR); only 8% of binding sites were located more than 10kb away from any protein coding gene. This strongly suggests a role for ING1 binding in the regulation of gene expression.

**Figure 1.**
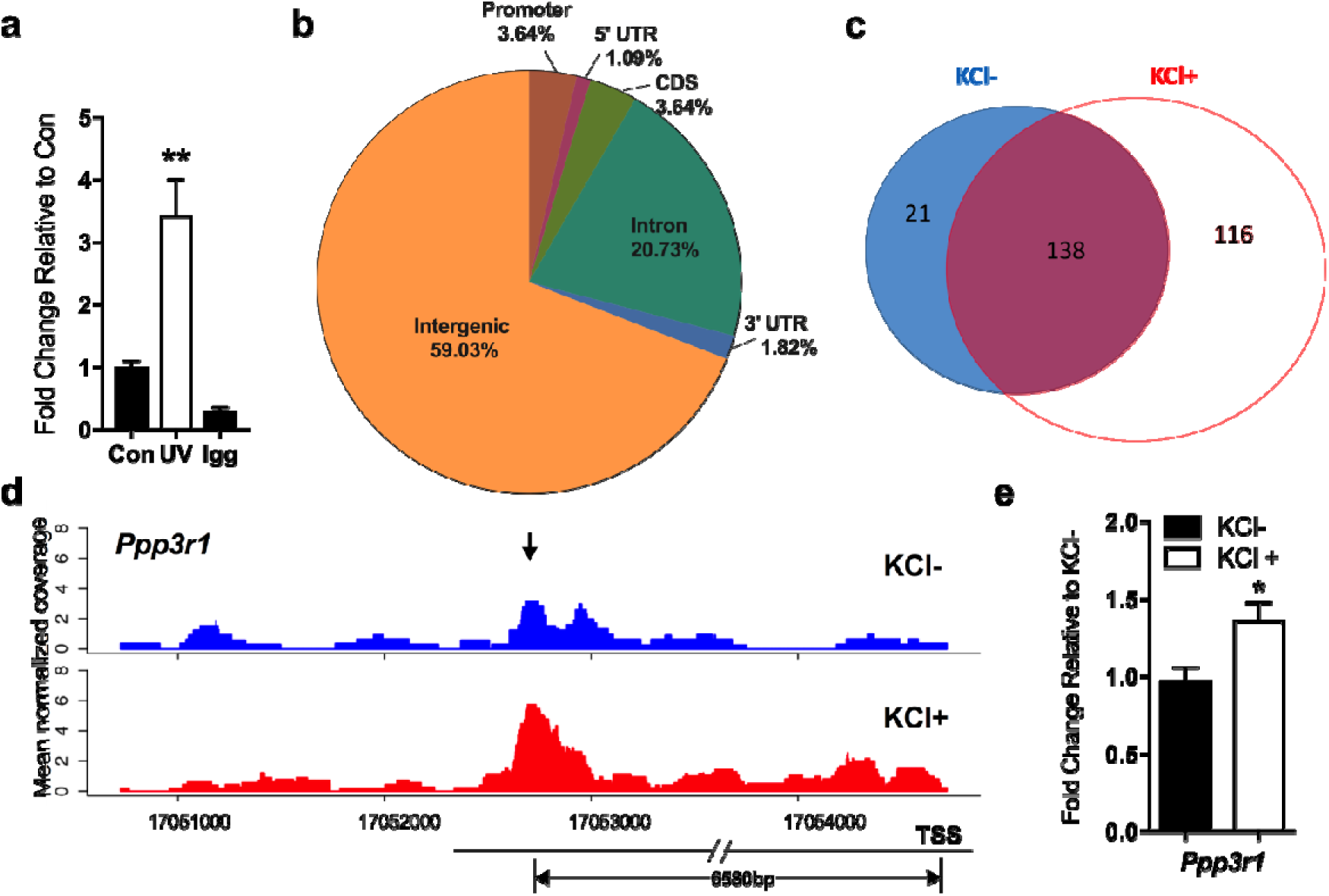
Genome-wide profiling reveals the dynamic landscape of ING1 binding sites in mouse primary cortical neurons following KCl-induced depolarisation. **(a)** The antibody detects accumulation of ING1 at the promoter of the known ING1 target *Mageb1* in UV-irradiated MEFs (n=4) compared to untreated MEFs (n=4) and was therefore considered suitable for ChIP. **(b)** Distribution analysis of ING1 ChIP-seq peaks identified in 3 hours KCl-stimulated (KCl+) primary cortical neurons, shown as a percentage of reads corresponding to each genomic region. **(c)** ING1 ChIP-seq peak counts in KCl− (n=3) and KCl+ (n=3) groups. Blue and red represent KCl− and KCl+, respectively. **(d)** Representative ING ChIP-seq traces for *Ppp3r1*, showing a consistent increase in ING1 binding in the KCl+ group (arrow). Blue and red traces represent immunoprecipitated coverages. The potential ING1 binding site is 6580bp upstream from the TSS of *Ppp3r1.* **(e)** ING1 ChIP-qPCR validation of ING1 binding to *Ppp3r1* under KCl depolarisation conditions (20mM, 3 h) (n=4-6, Student’s t-test, *p <0.05). Data represent mean ±SEM.

Among the 116 ING1 binding sites which were specific to the KCl+ group, we selected 5 for validation by ING1 ChIP-qPCR (Figure 1d, 1e and Figure 2). Figure 1d shows the mapping of ING1 ChIP-seq reads to a representative candidate gene, protein phosphatase 3 regulatory subunit B, alpha (*Ppp3r1*) for one KCl+/KCl− experimental pair. KCl-induced depolarisation led to an increase in ING1 binding to a region 6580bp upstream of the *Ppp3r1* TSS (upstream regulatory element/URE). The ING1 binding profiles for an additional four candidate genes *(Plcb4, Cdk19, Cacnb2* and *Lrrc7)* are shown in Figure 2; these binding profiles show that ING1 potentially interacts with intron, promoter, coding sequence (CDS) and intergenic regions separately. Taken together, these findings indicate that ING1 binding is activity-induced, and supports the hypothesis that ING1 is acting as a dynamic epigenetic regulator that is likely to be involved in the activity-dependent regulation of gene expression in primary cortical neurons.

**Figure 2.**
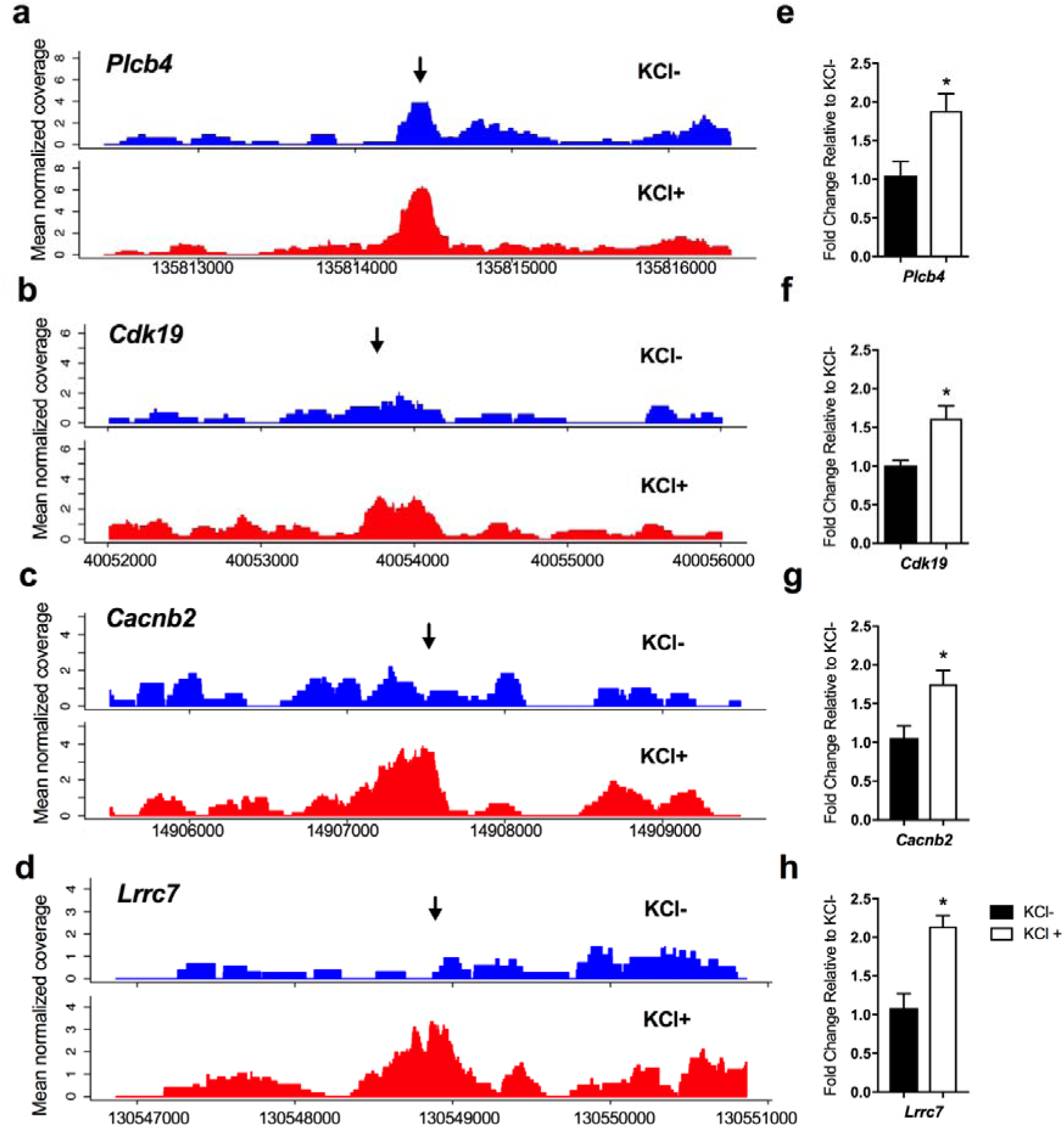
ING1 ChIP-seq plots of selected genes highlighting dynamic peaks in the presence (KCl+) or absence (KCl−) of KCl-induced depolarisation. Dynamic peaks were identified within the intron of *Plcb4* **(a)**, the promoter of *Cdk19* **(b)**, the CDS of *Cacnb2* **(c)**, and the intergenic region in front of *Lrrc7* **(d)** respectively; the arrows indicate the peak regions. Y-axis represents mean of normalised coverage. Increased ING1 occupancy at each locus was confirmed by ChIP-qPCR **(e-h)** under KCl depolarisation (20mM, 3h) (n=4-5, Student’s t-test, *p <0.05). Data represent mean ±SEM.

### Knockdown of *ING1* blocks induction of *Ppp3r1* by KCl stimulation

*Ppp3r1* encodes a regulatory subunit of calcineurin, a protein phosphatase that is involved in calcium signalling in neurons (Baumgärtel & Mansuy, 2012). Disruption of calcineurin activity has severe effects on memory in animal models (Malleret *et al.*, 2001; Zeng *et al.*, 2001; Cottrell *et al.*, 2013), making *Ppp3r1* an interesting candidate gene in the context of neuronal function. The binding of ING1 at the *Ppp3r1* locus in primary cortical neurons occurs at a site 6580bp upstream of the TSS. This KCl+ specific binding site is one of the predominantly intergenic ING1 binding sites identified by ChIP-Seq, and ING1 occupancy at this site shows an obvious increase in response to KCl stimulation, with higher normalised read coverage in KCl+ samples than that in KCl− samples (Figure 1d).

To determine the functional role of ING1 in the epigenetic regulation of gene expression, we designed lentiviral short-hairpin RNA (shRNA) constructs (Figure 3a) based on our previously published protocols (Lin *et al.*, 2011). Three different *Ing1* shRNAs (Table 1) were validated by Western blot and all produced a significant knockdown of ING1 at the protein level, compared to a universal shRNA control which has no specificity to any known mouse transcripts. Furthermore, treatment of primary cortical neurons with a cocktail of all 3 *Ing1* shRNAs blocked neural activity-induced ING1 binding at the *Ppp3r1* locus (Figure 3b).

**Figure 3.**
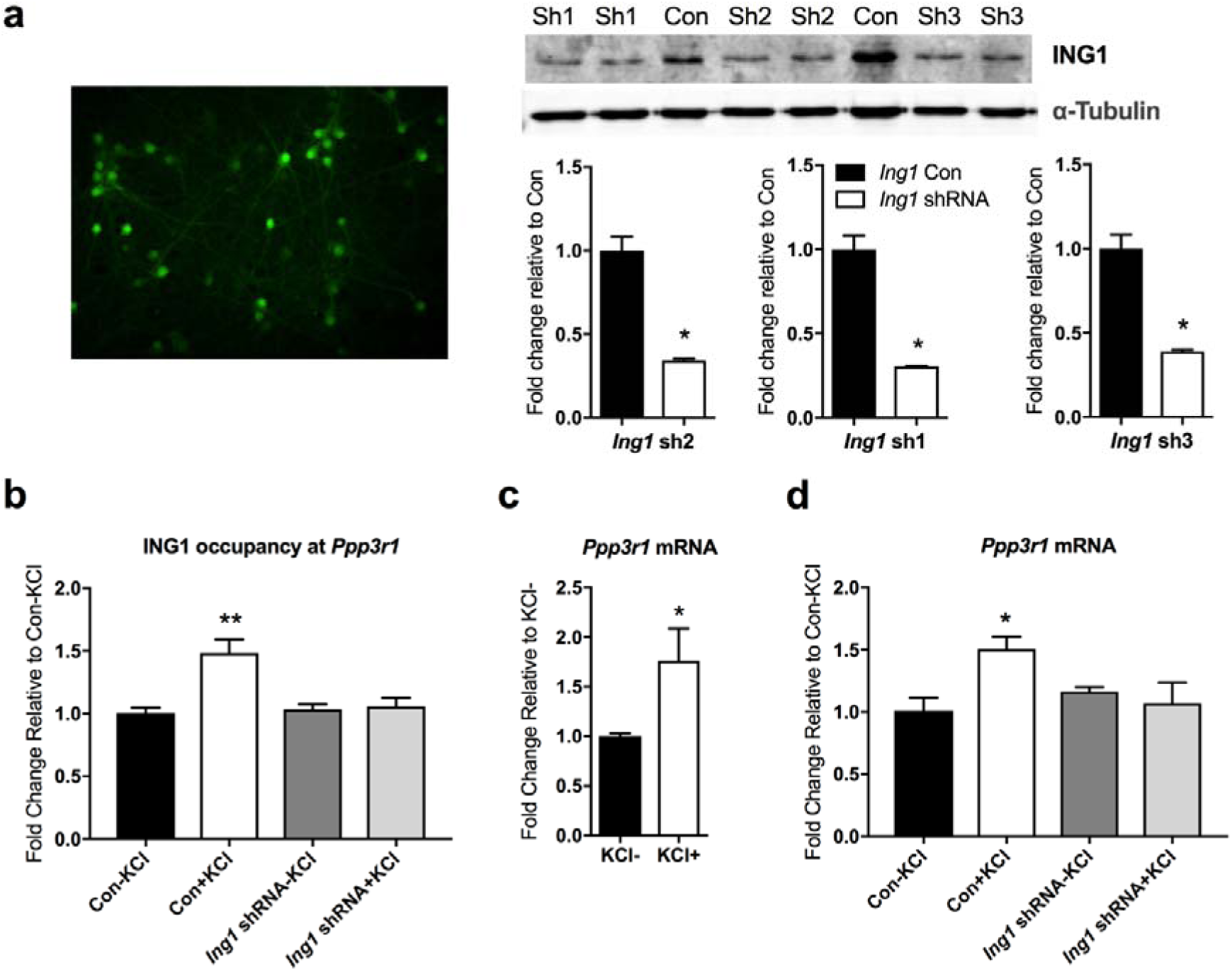
Activity-dependent ING1 occupancy is associated with increased *Ppp3r1* mRNA expression in primary cortical neurons. **(a)** shRNA targeting *Ing1* inhibits ING1 protein expression in primary cortical neurons. Left; representative image of neurons treated with shRNA-expressing lentivirus. Right; quantitative Western blot analysis confirms that each of the three *Ing1* shRNAs tested (sh1, sh2, sh3) knocks down expression of ING1 compared to the universal shRNA control (Con) (n=3, Student’s t-test, *p <0.05). **(b)** *Ing1* knockdown eliminated activity-induced ING1 occupancy at the *Ppp3r1* upstream regulatory region after 3 hours of KCl-induced depolarisation (n=3-4, one-way ANOVA, F (3, 12) = 10.36, **p<0.01). **(c)** *Ppp3r1* mRNA expression is inducible by KCl-induced depolarisation in primary cortical neurons (20mM_;_ 3h) (n=3-4, Student’s t-test, *p <0.05) **(d)** Expression of *Ppp3r1* mRNA is inhibited by *Ing1* knockdown (n=3-5, one-way ANOVA, F(3, 9) = 4.446, *p <0.05). ‘*Ing1* shRNA’ = cocktail of sh1, sh2 and sh3-expressing lentiviruses. Data represent mean ±SEM.

**Table 1.**
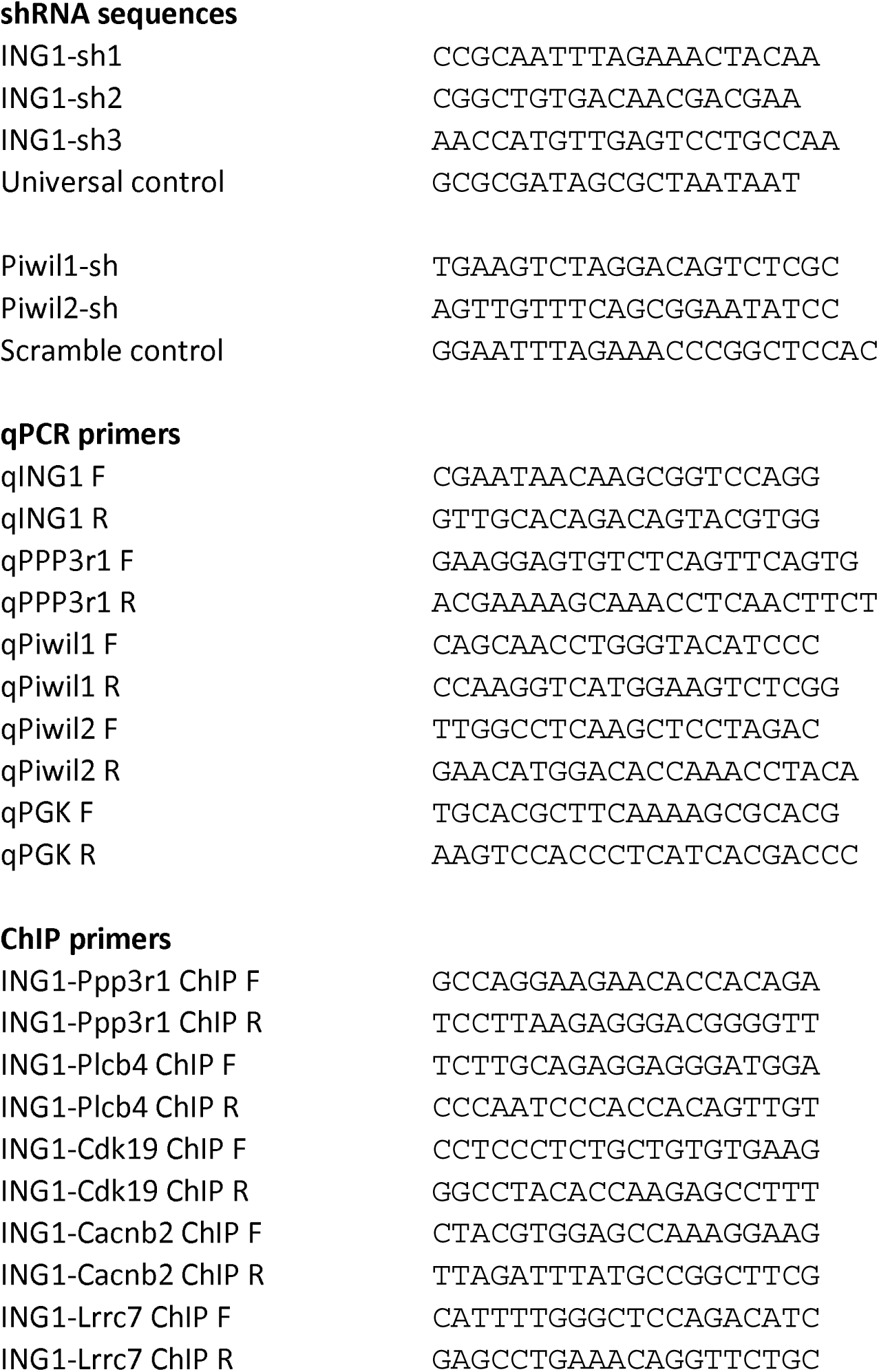
List of primer and shRNA sequences used in this study.

Stimulation of primary cortical neurons with KCl results in a robust upregulation of *Ppp3r1* mRNA, which reached approximately 160% of its baseline expression (Figure 3c). In neurons treated with lentivirus carrying shRNAs targeting *Ing1*, the induction of *Ppp3r1* by KCl-induced depolarisation was entirely abolished, revealing the importance of ING1 deposition at the *Ppp3r1* locus for the activity-dependent induction of the *Ppp3r1* gene (Figure 3d). Taken together, these observations indicate that the KCl-mediated increase in *Ppp3r1* mRNA expression is due, at least in part, to increased ING1 binding at the *Ppp3r1* URE region.

### Piwi proteins are expressed in primary cortical neurons

As part of our exploration of the crosstalk between potential epigenetic regulatory mechanisms and neuronal gene expression, we also examined the expression of the Piwi-like genes *Piwil1* and *Piwil2* in primary cortical neurons. We found that *Piwil1* and *Piwil2* are both expressed in primary cortical neurons (Figure 4a, b). This is consistent with the observation of neuronal expression of piRNAs in several previously published studies (Lee *et al.*, 2011; Saxena *et al.*, 2012; Ghosheh *et al.*, 2016; Nandi *et al.*, 2016; Roy *et al.*, 2017). In addition, we found that *Piwil1* and *Piwil2* are both induced by KCl-induced depolarisation. After 3 hours of KCl exposure, *Piwil1* was upregulated threefold, while *Piwil2* expression doubled (Figure 3a, b). These data indicate the presence of low-level, but potentially biologically relevant, expression of components of the Piwi pathway in mouse neurons, and support a link between the Piwi pathway and neuronal activity.

### Knockdown of *Piwil1* and *Piwil2* blocks *Ppp3r1* mRNA expression and prevents ING1 binding to the *Ppp3r1* upstream regulatory region

Given the evidence that the Piwi pathway lies upstream of DNA repair, in which ING1 is involved as an epigenetic regulator, we next asked whether the effect of disruption of the Piwi pathway on *Ppp3r1* expression was similar to that of *Ing1* knockdown. Given that little is known about the function of either *Piwil1* or *Piwil2* in neurons, we chose to investigate the function of the pathway as a whole by simultaneously knocking down both *Piwil1* and *Piwil2.* We used adeno-associated viral vectors (AAVs) carrying shRNA sequences directed against *Piwil1, Piwil2*, and as a negative control, a scrambled sequence with no targets in the mouse genome (Table 1.) We confirmed that AAVs carrying shRNA against *Piwil1* and *Piwil2* significantly knocked down expression of target mRNAs in primary cortical neurons four days after application (Figure 4c, 4d.)

**Figure 4.**
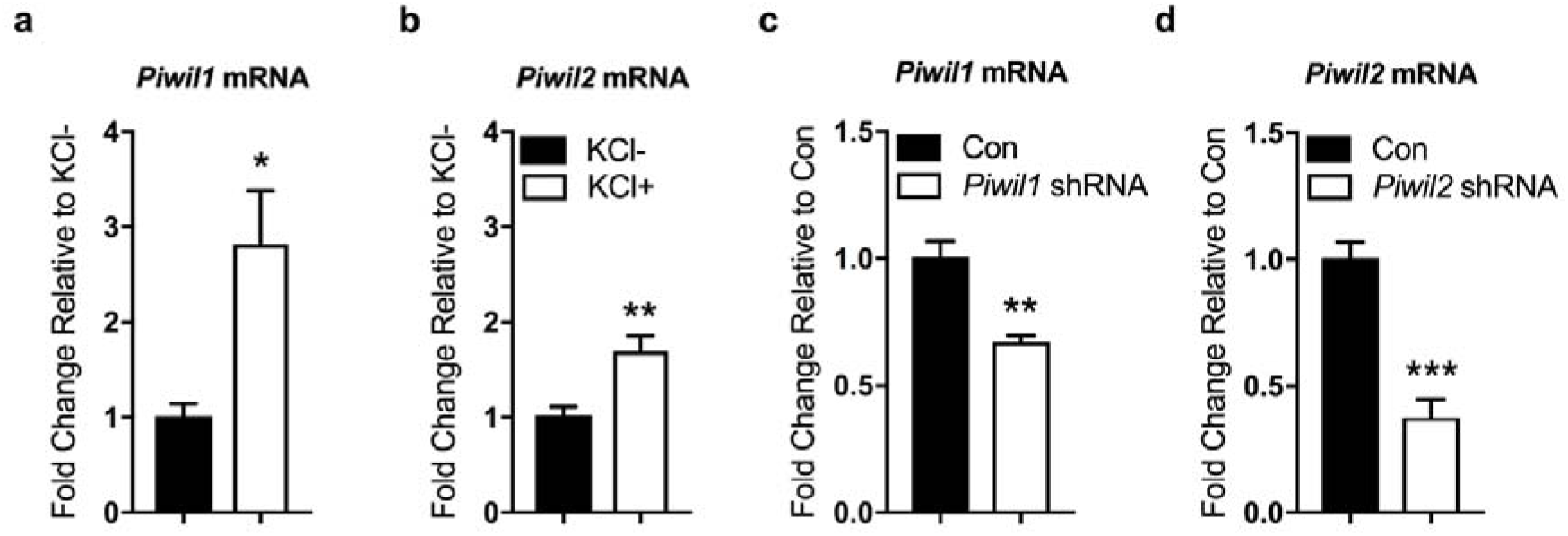
The expression of *Piwil1* and *Piwil2* are induced by KCl in primary cortical neurons. *Piwil1* **(a)** and *Piwil2* **(b)** mRNA is detected in primary cortical neurons at baseline; mRNA of both *Piwil1* and *Piwil2* significantly increases following 3 hours of KCl stimulation (n=4-6, Student’s t-test, *p<0.05, **p <0.01). Expression of *Piwil1* **(c)** and *Piwil2* **(d)** is significantly reduced in primary cortical neurons 4 days after treatment with the respective shRNA-expressing AAVs, compared to neurons treated with a non-targeting control. (n=4, Student’s t-test, **p <0.01, ***p <0.001).

We next examined the expression of *Ppp3r1* mRNA in primary cortical neurons treated with both *Piwil1-*shRNA and *Piwil2*-shRNA AAVs. Simultaneous knockdown of *Piwil1* and *Piwil2* blocked the neural activity-induced upregulation of *Ppp3r1* mRNA expression (Figure 5a). Surprisingly, we also found that disruption of the Piwi pathway abolishes the activity-dependent *Ppp3r1* induction by modulating ING1 binding dynamics. ChIP-qPCR revealed that knockdown of *Piwil1* and *Piwil2* eliminated the KCl-dependent increase in ING1 occupancy at the *Ppp3r1* URE (Figure 5b), thereby blocking the ING1-mediated upregulation of this gene in response to neuronal activation.

**Figure 5.**
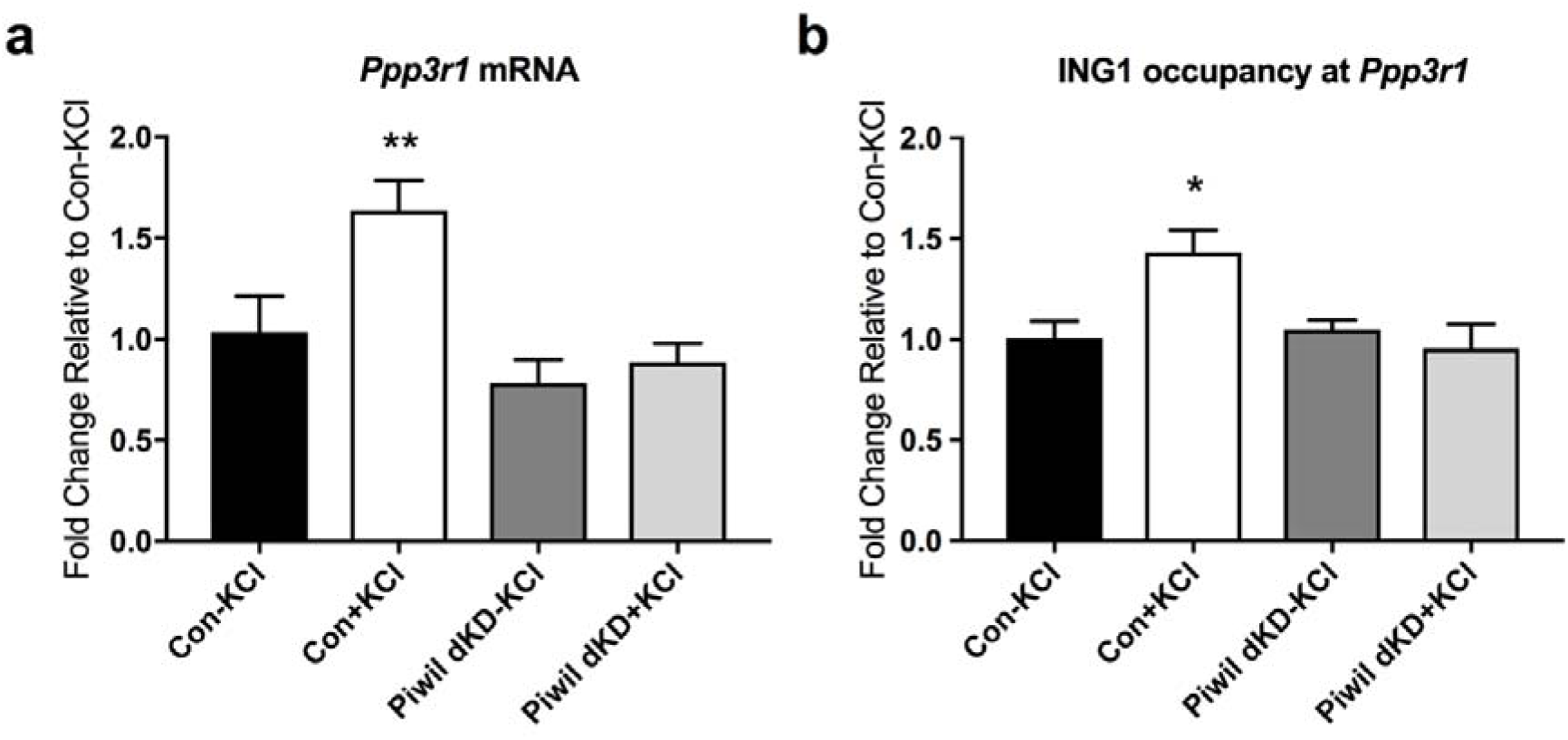
Regulation of Ppp3r1 mRNA expression by ING1 is mediated by the Piwi pathway. **(a)** *Ppp3r1* mRNA induction by KCl-induced depolarisation (20mM, 3h) in primary cortical neurons is inhibited by simultaneous knockdown of *Piwil1* and *Piwil2* (n=3-5, one way ANOVA, F (3, 10) = 8.101, **p <0.01). **(b)** Simultaneous knockdown of *Piwil1* and *Piwil2* eliminates activity-induced ING1 occupancy at the *Ppp3r1* upstream regulatory region under KCl-induced depolarisation (n=3-4, one way ANOVA, F (3, 9) = 4.547, *p <0.05). Con = scrambled non-targeting shRNA. Data represent mean ±SEM.

### Activating histone marks accumulate at the *Ppp3r1* locus in response to neuronal activation

After discovering the activity-induced increase in ING1 binding at the *Ppp3r1* URE, we next looked for evidence of change in the local chromatin landscape that is associated with ING1 binding and *Ppp3r1* mRNA expression. Given the previously identified interaction between ING1 and histone modification (Skowyra *et al.*, 2001; Feng *et al.*, 2002; Binda *et al.*, 2008), we reasoned that ING1 may interact with the local chromatin landscape in an activity-induced manner, thereby providing a further link between ING1 binding and the epigenetic regulation of gene expression. To explore this possibility, we used ChIP-qPCR to interrogate the chromatin environment surrounding the ING1 binding site upstream of the *Ppp3r1* gene for increases in four permissive histone marks: H3K4me3, H3K4me1, H3K9ac and H3K14ac.

We found that there was no effect of KCl stimulation on the occupancy of H3K4me1 or H3K14ac at the *Ppp3r1* URE; these two histone marks are typically present within active enhancer regions (Wang *et al.*, 2008; Heintzman *et al.*, 2009). However, ChIP-qPCR revealed increased occupancy (>2.5-fold) of H3K4me3 at the *Ppp3r1* URE after 3 hours of KCl stimulation (Figure 6a). H3K4me3 is primarily associated with active promoters and supports the initiation of transcription (Guenther *et al.*, 2007; Wang *et al.*, 2008), but also occurs within active enhancer regions (Chen *et al.*, 2015). Additionally, we detected approximately 1.5-fold enrichment of H3K9ac at the *Ppp3r1* URE in the KCl-stimulated cells (Figure 6b); this histone mark is broadly associated with active promoters (Wang *et al.*, 2008; Heintzman *et al.*, 2009) and is associated with euchromatin at enhancers and other sites (Schebesta *et al.*, 2007). The increase in occupancy of the activating marks H3K4me3 and H3K9ac at the *Ppp3r1* URE following KCl stimulation indicates that the chromatin state at this site has become more permissive for transcription, and for binding of transcription factors and regulatory elements. This finding is consistent with both the increased ING1 binding and enhanced *Ppp3r1* expression observed under depolarising conditions.

**Figure 6.**
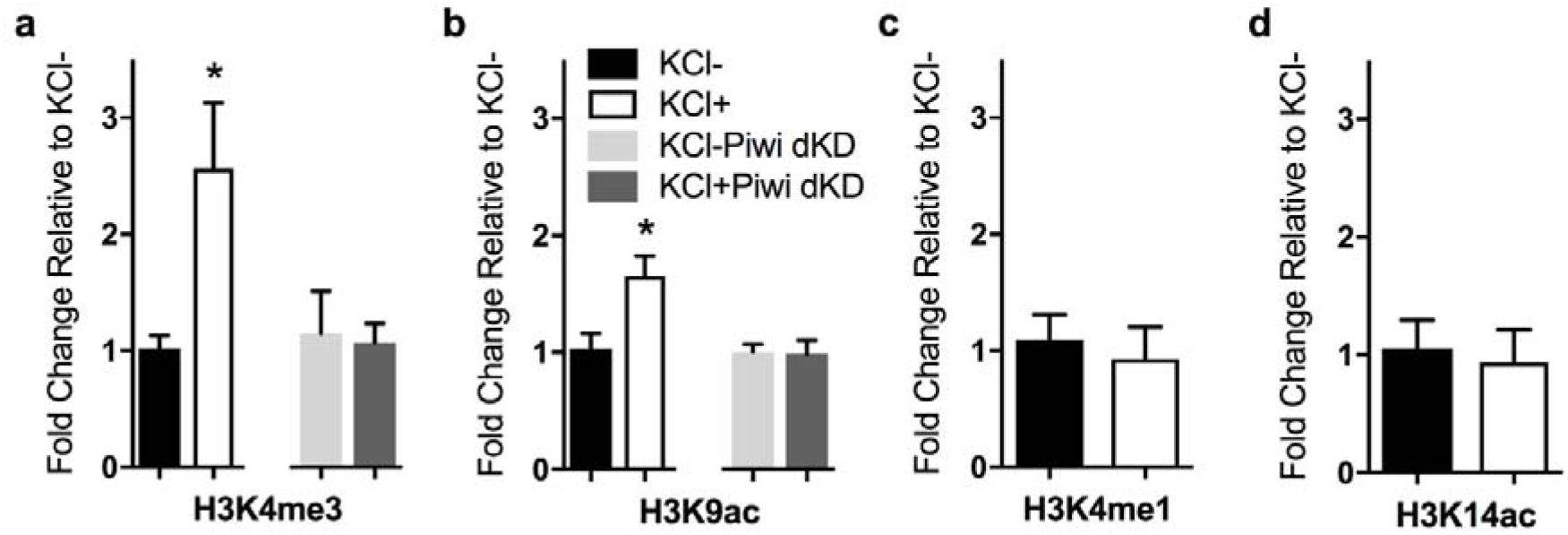
ING1 binding at the Ppp3r1 locus is associated with activating histone marks, which are abolished following knockdown of *Piwil1* and *Piwil2.* There is a significant increase in H3K3me3 **(a)** and H3K9ac **(b)** occupancy at the *Ppp3r1* upstream regulatory region under KCl-induced depolarisation(20mM, 3h), but no significant difference in H3K4me1 **(c)** and H3K14ac **(d)** occupancy. Simultaneous knockdown of *Piwil1* and *Piwil2* prevents accumulation of both H3K4me3 **(a)** and H3K9ac **(b)** at this locus. (n=3-4, Student’s t-test, *p <0.05). Data represent mean ±SEM.

We next asked whether disruption of the Piwi pathway, which is known to have a role in chromatin remodelling, could perturb the development of this permissive chromatin state in response to neuronal activity. Interestingly, we observed that simultaneous knockdown of Piwil2 prevented accumulation of either H3K4me3 or H3K9ac at the *Ppp3r1* URE in response to KCl stimulation (Figure 6a, 6b), indicating that the Piwi pathway is upstream of histone modification at the *Ppp3r1* locus in this system.

## Discussion

In this study, we have demonstrated that 1) ING1, a protein with a wide range of epigenetic and regulatory functions, exhibits altered binding dynamics in response to neuronal activity in primary cortical neurons *in vitro*, 2) increased occupancy of ING1 at an upstream regulatory region near the *Ppp3r1* gene is necessary for the activity-induced upregulation of *Ppp3r1* mRNA expression, and 3) disruption of the Piwi pathway blocks the upregulation of *Ppp3r1* by preventing ING1 binding at the *Ppp3r1* URE. ING1 is a multifaceted epigenetic regulator, and many of the epigenetic processes to which it contributes are known to be important for neuronal function. However, our results represent the first specific evidence that ING1 is involved in regulating activity-dependent gene expression in normal neurons.

Consistent with its role as a tumour suppressor, *Ing1* is a DNA damage response gene (Cheung Jr *et al.*, 2000; Cheung *et al.*, 2001; Ceruti *et al.*, 2013) that is involved in DNA repair through several pathways (Ceruti *et al.*, 2013). It has recently been reported that DNA double-strand breaks occur in neurons in an experience-dependent manner (Suberbielle *et al.*, 2013; Bellesi *et al.*, 2016), and that DNA breaks occur at the promoters of neuronal early-response genes (Madabhushi *et al.*, 2015) and other stimulus-inducible genes (Bunch *et al.*, 2015) and are required for their transcription. These findings raise the possibility that proteins involved in the recognition of DNA damage and subsequent DNA repair could act as a mechanism to target epigenetic or transcriptional machinery to loci in response to experience, such as learning. This possibility led us to ask whether disruption of the Piwi pathway affects activity-dependent gene regulation in a similar way to knockdown of *Ing1*. In addition to the mounting evidence that the Piwi pathway is functional in neurons, other unexpected roles for this pathway in a range of somatic cell types have recently been revealed (Ross *et al.*, 2014). For example, Yin *et al.* showed that *Piwil2*, which is normally silenced in fibroblasts, is transiently activated in response to DNA damage (Yin *et al.*, 2011). This is normally followed by histone H3 acetylation (H3K9ac and H3K14ac) which results in chromatin relaxation, allowing DNA repair enzymes to access the DNA. In cells deficient in PIWIL2, histone acetylation and chromatin relaxation do not occur in response to DNA damage, resulting in defective DNA repair and revealing that PIWIL2 is necessary at the first stage of the DNA damage response.

We speculate that the Piwi pathway is upstream of ING1 during activity-dependent regulation of *Ppp3r1* in primary cortical neurons. Specifically, it is known that DNA breakage occurs in response to neuronal activity (Suberbielle *et al.*, 2013; Bellesi *et al.*, 2016) and Piwi proteins are known to respond to DNA breakage by promoting chromatin relaxation (Yin *et al.*, 2011). We observed that simultaneous knockdown of *Piwil1* and *Piwil2* prevents the accumulation of the permissive histone marks H3K4me3 and H3K9ac at the *Ppp3r1* URE in response to KCl stimulation. One of these histone marks, H3K9ac, was previously found to increase in response to DNA damage in fibroblasts; this increase did not occur in PIWIL2^-/-^ cells (Yin *et al.*, 2011). Taken together, these observations strongly suggest that knockdown of *Piwil1* and *Piwil2* results in inappropriate chromatin structure which impedes access of ING1 at the *Ppp3r1* promoter, resulting in the loss of both ING1 binding and *Ppp3r1* induction that is observed in the *Piwil1/Piwil2* knockdown neurons. The Piwi pathway is primarily involved in transcriptional and epigenetic silencing, and it is not currently clear how Piwi-like proteins affect histone modifications. However, the Piwi pathway is known to be involved in chromatin remodelling in *Drosophila* (Pal-Bhadra *et al.*, 2004; Brower-Toland *et al.*, 2007; Huang *et al.*, 2013), including one case where a specific piRNA drives a euchromatic state at a complementary locus (Yin & Lin, 2007).

It is also possible that *Piwil1/Piwil2* knockdown has an effect on ING1-mediated *Ppp3r1* expression through a less direct mechanism. Given that H3K4me3 is able to recruit ING1, it is possible that ING1 binding at the *Ppp3r1* URE is driven by the accumulation of this epigenetic mark. If this is the case, any activity of the Piwi pathway which promotes accumulation of H3K4me3 at the *Ppp3r1* URE could contribute to the loss of ING1 binding observed in neurons where *Piwil1* and *Piwil2* are knocked down. Several studies have detected expression of piRNAs in neurons and observed altered piRNA expression in response to various stimuli (Lee *et al.*, 2011; Rajasethupathy *et al.*, 2012; Saxena *et al.*, 2012; Ghosheh *et al.*, 2016; Nandi *et al.*, 2016). The targets of virtually all of these small regulatory RNAs remain unidentified, but could certainly include chromatin-modifying enzymes, and other genes which may affect ING1 binding or *Ppp3r1* expression in other ways. Another possibility is that *Piwil1* and/or *Piwil2* may be acting independently of piRNAs; this has been suggested by Viljetic *et al.* who found that although *Piwil1* knockdown disrupts mouse corticogenesis, it has little effect on cortical piRNA expression (Viljetic *et al.*, 2017).

The downstream target identified by this study, *Ppp3r1*, encodes a regulatory subunit of calcineurin, a calcium and calmodulin-dependent serine/threonine protein phosphatase. Calcineurin is one of the most abundant protein phosphatases in the brain, and modulates the function of other proteins in response to calcium signalling (Baumgärtel & Mansuy, 2012). Calcineurin has been extensively implicated in learning and memory (Malleret *et al.*, 2001; Zeng *et al.*, 2001; Lin *et al.*, 2003; Havekes *et al.*, 2006; Baumgärtel & Mansuy, 2012; Cottrell *et al.*, 2013), schizophrenia (Gerber *et al.*, 2003; Miyakawa *et al.*, 2003; Eastwood *et al.*, 2005) and depression (Seimandi *et al.*, 2013; Yu *et al.*, 2013). Altered expression or regulation of one of its subunits, as we observed in the present study, could therefore be expected to have broad implications for neuronal function. Calcineurin has also been implicated in Alzheimer’s disease due to its effects on Tau metabolism (Gong *et al.*, 1994; Luo *et al.*, 2008; Karch *et al.*, 2013), with a genetic variant of *Ppp3r1* being strongly associated with the rapid progression of Alzheimer’s disease in humans (Cruchaga *et al.*, 2010; Peterson *et al.*, 2014). This provides strong justification for further investigation of *Ppp3r1* as a gene of potential relevance to learning, memory and human health.

In conclusion, we report that ING1 is dynamically regulated in neurons and is involved in the epigenetic regulation of neuronal gene expression. We also show that ING1 binding to at least one target regulatory region, a site upstream of the TSS of *Ppp3r1*, depends on neuronal Piwi-like proteins. These findings provide important new insight into the epigenetic regulation of activity-induced gene expression in neurons, which has relevance for understanding neural plasticity, learning and memory.

## Acknowledgements

This work was supported by grants from the National Institutes of Mental Health (1R01MH109588 to T.W.B.) and Australian National Health and Medical Research Council (GNT1062570 to T.W.B.) and postgraduate scholarships from the Westpac Bicentennial Foundation and the Australian Government Research Training Program (administered by The University of Queensland) to L.J.L. We thank Ms. Rowan Tweedale for helpful editing of this paper.

